# Sixteen isotropic 3D fluorescence live imaging datasets of *Tribolium castaneum* gastrulation

**DOI:** 10.64898/2026.01.16.699888

**Authors:** Franziska Krämer, Stefan Münster, Frederic Strobl

## Abstract

Gastrulation is a pivotal phase of early embryogenesis during which initially uniform cells undergo coordinated movements and fate specification to establish the basic body plan. Although previous studies have advanced our understanding of gastrulation in the red flour beetle *Tribolium castaneum*, quantitative morphogenetic data remain limited. Here, we present the second Systematic Live Imaging Collection of Embryogenesis (SLICE-2), consisting of sixteen isotropic 3D fluorescence live imaging datasets that document *Tribolium* gastrulation and early germband elongation. Imaging was performed using light sheet fluorescence microscopy in conjunction with a homozygous transgenic line that expresses mEmerald-labeled nanobodies against histone H2A/H2B. Embryos were recorded along four orientations, and the resulting axial image stacks were subjected to multiview fusion to obtain isotropic 3D images of entire embryos with reduced shadowing artifacts. Datasets are provided as weighted-average fusion and fusion-deconvolution derivatives, and nuclei segmentation data are available for selected datasets and stages. SLICE-2 expands the resource arsenal for *Tribolium*, enabling comprehensive analyses of gastrulation dynamics and providing benchmark data for image processing, segmentation, and modeling approaches.

## Background & Summary

Understanding how a single cell gives rise to a complex, patterned multicellular organism is a central goal of developmental biology^1^. Gastrulation, a defining event of early embryogenesis, marks a phase of profound cellular reorganization in which initially uniform populations begin to move, fold, and adopt distinct identities. It results in the formation of the three germ layers—ectoderm, mesoderm, and endoderm—from which all embryonic tissues ultimately emerge^2^. Despite its central role, the mechanisms and dynamics of gastrulation exhibit striking diversity across the metazoan lineage^3,4^.

The red flour beetle *Tribolium castaneum* has become a well-established model organism in evolutionary developmental biology^5^. It combines the practical advantages of a short generation time and genetic tractability with a developmental mode considered more representative of ancestral insects than that of the fruit fly *Drosophila melanogaster*^6^. Previous work has provided important insights into diverse aspects of *Tribolium* gastrulation, including mesoderm invagination^7–9^, segment identity specification^10–15^, and extra-embryonic membrane (EEM) formation^13,16–19^ with more recent efforts focusing on biomechanical mechanisms^20,21^. Unlike *Drosophila*, which gives rise to a single EEM, the amnioserosa^22^, *Tribolium* forms two distinct extra-embryonic layers, the amnion and the serosa. During gastrulation, the serosa moves over the yolk and the condensing embryonic rudiment and eventually closes at an anterior-ventral position, enveloping all underlying structures throughout germband elongation and retraction before eventually rupturing and withdrawing at the beginning of dorsal closure^13,21,23^. Despite the biological significance of these processes, 3D live imaging data documenting the morphogenetic dynamics of *Tribolium* gastrulation remain scarce, and many aspects lack quantitative context. For example, the total numbers of blastodermal nuclei—and consequently the fractions that give rise to either embryonic tissue or EEMs—have never been determined and are currently available only as rough estimates^24^.

In this data descriptor, we present the second Systematic Live Imaging Collection of Embryogenesis (SLICE-2) consisting of sixteen isotropic 3D fluorescence live imaging datasets of *Tribolium castaneum* gastrulation. Data were acquired using light sheet fluorescence microscopy in conjunction with the AGOC{ATub’H2A/H2B°NB-mEmerald} #1 transgenic subline^25^, which expresses mEmerald-labeled nanobodies against histone H2A/H2B under control of the consecutively and ubiquitously active *tubulin alpha 1-like protein* promoter^26^. Imaging began at the end of blastoderm formation and continued for ∼24 h, covering gastrulation and part of germband elongation. All embryos hatched morphologically intact, all but one larva developed into healthy adults, and all but two of the remaining adults produced progeny.

Each embryo was recorded along four orientations spaced 90° apart, and the resulting image data were subjected to multiview fusion to obtain enhanced representations of the data with strongly reduced shadowing artifacts^27^ (Figure 1). The fused images were rotated around all three spatial axes to align embryonic and image axes, cropped to standardized sizes, and adjusted in brightness and contrast. Resulting isotropic 3D images are available as two different derivatives, either as weighted-average fusions, or as fusion-deconvolutions. Based on the fusion-deconvolution 3D images, the superficial blastodermal nuclei at the end of blastoderm formation as well as the serosa nuclei at the beginning of germband elongation were segmented in selected datasets, yielding averages (and standard deviations) of 3536 (± 99) and 848 (± 128) nuclei, respectively.

**Figure 1.**
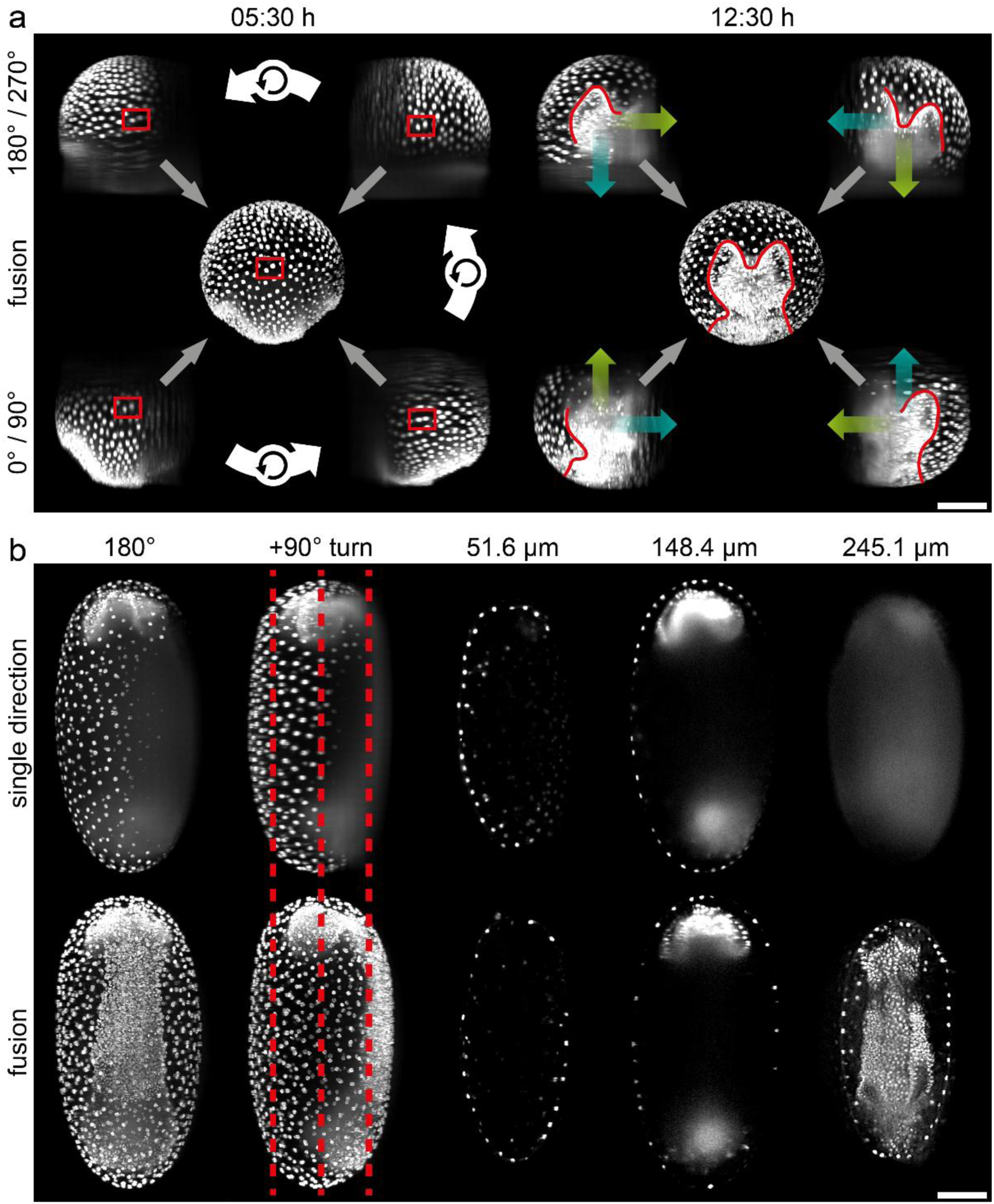
Multi-view fusion of imaging data obtained from a *Tribolium* embryo expressing mEmerald-labeled nanobodies against histone H2A/H2B. **(a)** Maximum projections along *y* of *z* stacks acquired along four directions spaced 90° apart (outer panels) and the corresponding fused 3D images (inner panels, hinted by gray arrows) at two time points from the same dataset. At 05:30 h, white arrows indicate the acquisition order of the four directions, which were rotated back in silico by 90° to ensure that all panels display the embryo from the same perspective. Red rectangles highlight the same two nuclei across panels. At 12:30 h, cyan and green arrows denote the illumination and detection axes, respectively. Red outlines indicate the part of the developing germband that is clearly visible in the respective projections, i.e. not obscured by shadowing artifacts. The full contour becomes discernible only in the fused image. **(b)** Maximum projections along *z* (first column) and *x* (second column), and optical sections (third to fifth column) of two corresponding 3D images, a ‘single direction’ version derived from a *z* stack with interplane interpolation, and a ‘fusion’ version representing the image obtained at the end of the image processing procedure. Red dashed lines mark the positions of the optical sections. The reduction of shadowing artifacts and consequently, the improvement in signal quality, is most apparent at a depth of ∼245 µm. Image data from dataset DS0005. Scale bars, 100 µm.

SLICE-2 complements existing image data resources on *Tribolium* gastrulation (as cited above) by filling an important gap: since the fusion derivatives exhibit comparable quality around the entire embryo, they enable statistically robust, global assessments of the morphogenetic processes underlying gastrulation. In addition, the non-fused *z* stacks and fused 3D images provide benchmark material for evaluating and comparing alternative fusion and/or deconvolution workflows, while the accompanying segmentation data (exemplified in Supplementary Movies 1 and 2) can be directly used for downstream analyses and modeling or serve as ground truth for further manual, semi-automatic, or automated segmentation approaches.

## Methods

### *Tribolium castaneum* transgenic lines and rearing

Imaging cultures of the transgenic AGOC{ATub’H2A/H2B°NB-mEmerald} #1 *Tribolium castaneum* subline^25^, which expresses mEmerald-labeled nanobodies against histone H2A/H2B, were maintained in populations of about 200–500. Cultures were reared on growth medium (full grain wheat flour (SP061036, Demeter) supplemented with 5% (wt/wt) inactive dry yeast (62-106, Flystuff)) in one-liter glass bottles in a 12:00 h light / 12:00 h darkness cycle at 32°C and 70% relative humidity (DR-36VL, Percival Scientific). All animal-related experiments were approved by the institutional ethics committee of the Goethe University (Tierschutzkommission der Goethe-Universität, file number BB-20250515-St) and conducted in accordance with the German Animal Welfare Act (Tierschutzgesetz/Tierschutz-Versuchstierordnung, based on ETS No.123^28^ and EU Directive 2010/63/EU^29^) as well as the ARRIVE 2.0 guidelines^30^.

### Light sheet fluorescence microscopy

Light sheet fluorescence microscopy was performed using a sample chamber-based digital scanned laser light sheet fluorescence microscope featuring single-sided illumination and single-sided detection (DSLM)^31^. A dynamic light sheet is generated by rapidly scanning a collimated Gaussian laser beam with a two-axes piezo-driven scanning mirror (M-116.DG, Physik Instrumente GmbH & Co KG). Illumination was provided by a 488 nm / 20 mW diode laser (PhoxX 488-20, Omicron Laserprodukte GmbH) with a 488 nm cleanup filter (xX.F488, Omicron Laserprodukte GmbH). Excitation light was delivered through a 2.5× NA 0.06 EC Epiplan-Neofluar objective (422320-9900-000, Carl Zeiss AG) while emission light was collected through a 10× NA 0.3 W N-Achroplan objective (420947-9900-000, Carl Zeiss AG). Detection was achieved using a 525/50 single-band bandpass filter (FF03-525/50-25, Semrock/AHF Analysentechnik AG) and a 16-bit (effectively 14-bit) high-resolution charge-coupled device camera (Clara, Andor). Conventionally^32^, the illumination axis is defined as *x*, the rotation axis as *y*, and the detection axis as *z*, with *y* aligned with the direction of gravity. Three micro-translation stages (M-111.2DG, Physik Instrumente GmbH & Co KG) were used for sample translation along *x*, *y*, and *z*, and one precision rotation stage (M-116.DG, Physik Instrumente GmbH & Co KG) for rotation around *y*. The effective laser power was routinely monitored with an optical power and wavelength meter (OMM-6810B and OMH-6703B, Newport).

### Embryo collection, preparation, mounting, live imaging and retrieval

Embryo collection and preparation were performed as described previously^33,34^. After collection, embryos were incubated at room temperature (23°C ± 1°C) for 15 h to reach the end of blastoderm formation. Embryos were mounted using the cobweb holder method, and an agarose column (∼460 µm diameter) containing a 5% (v/v) suspension of 1 µm fluorescent microspheres (T7282, Thermo Fisher Scientific) was positioned 50–150 µm below each embryo. In the DSLM, axial image (*z*) stacks of embryos were recorded together with a portion of the fluorescent microspheres-containing agarose column in one fluorescence channel along four directions in 90° steps and at up to 51 time points with an interval of 30 min (total recording duration up to 25 h). Embryos and microspheres were illuminated with a laser power of 135 µW during a 50 ms exposure time window of the camera. All *z* stacks had a lateral voxel pitch of 0.645 µm and an axial voxel pitch of 2.58 µm (i.e. a lateral-to-axial pitch ratio of 1:4). After imaging, embryos were retrieved from the microscope chamber as described previously^27^.

### Image processing

All sixteen datasets (Table 1) were processed following an identical workflow. Image processing started with the basic branch (denoted with ‘B’ in subfolder names), which encompasses a simple processing routine for quick visual inspection as described previously^25^, except that no histogram matching was performed. Further image processing was performed exclusively using Fiji^35^ (based on ImageJ^36^ Version 1.53f).

**Table 1.**
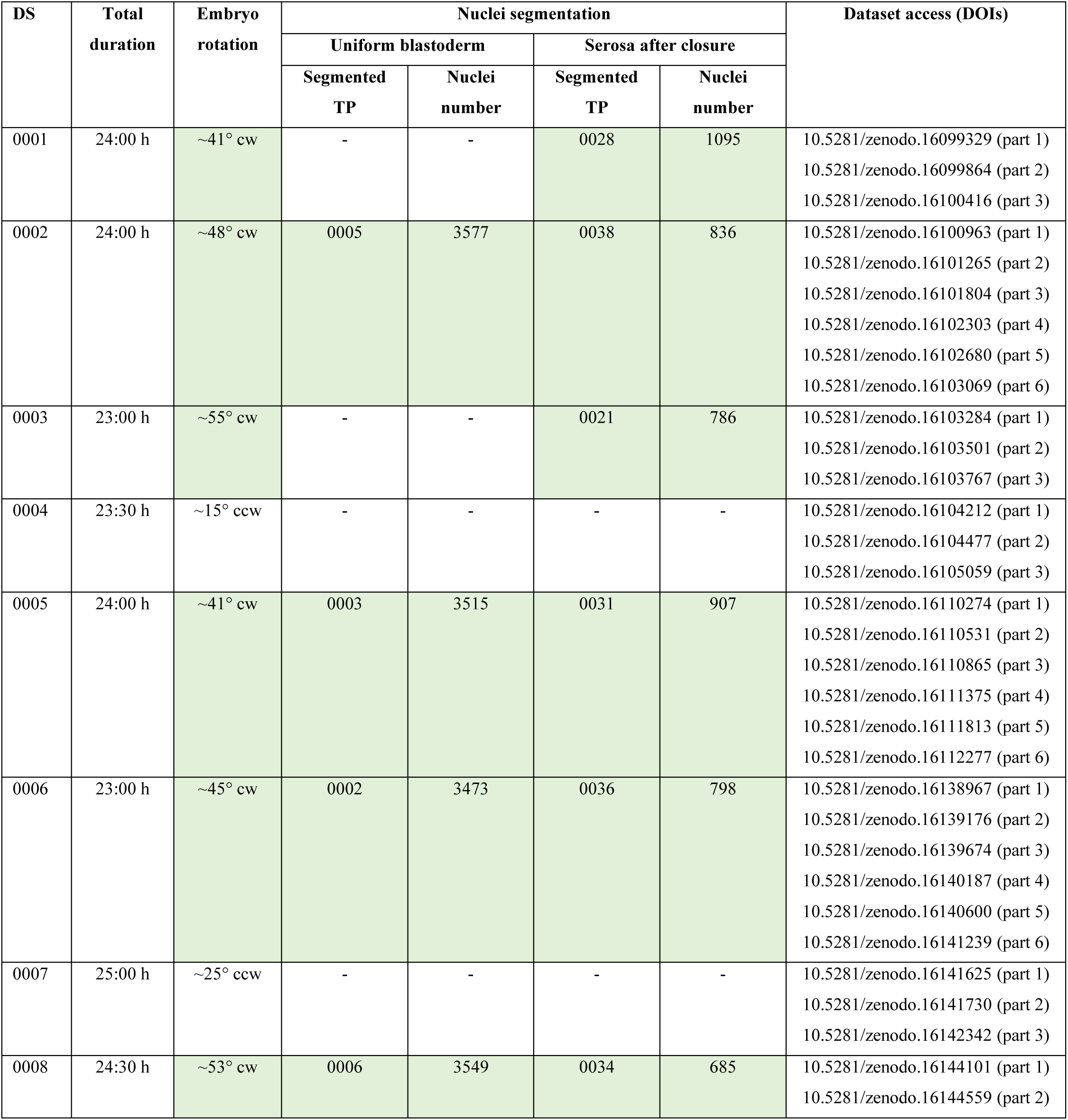
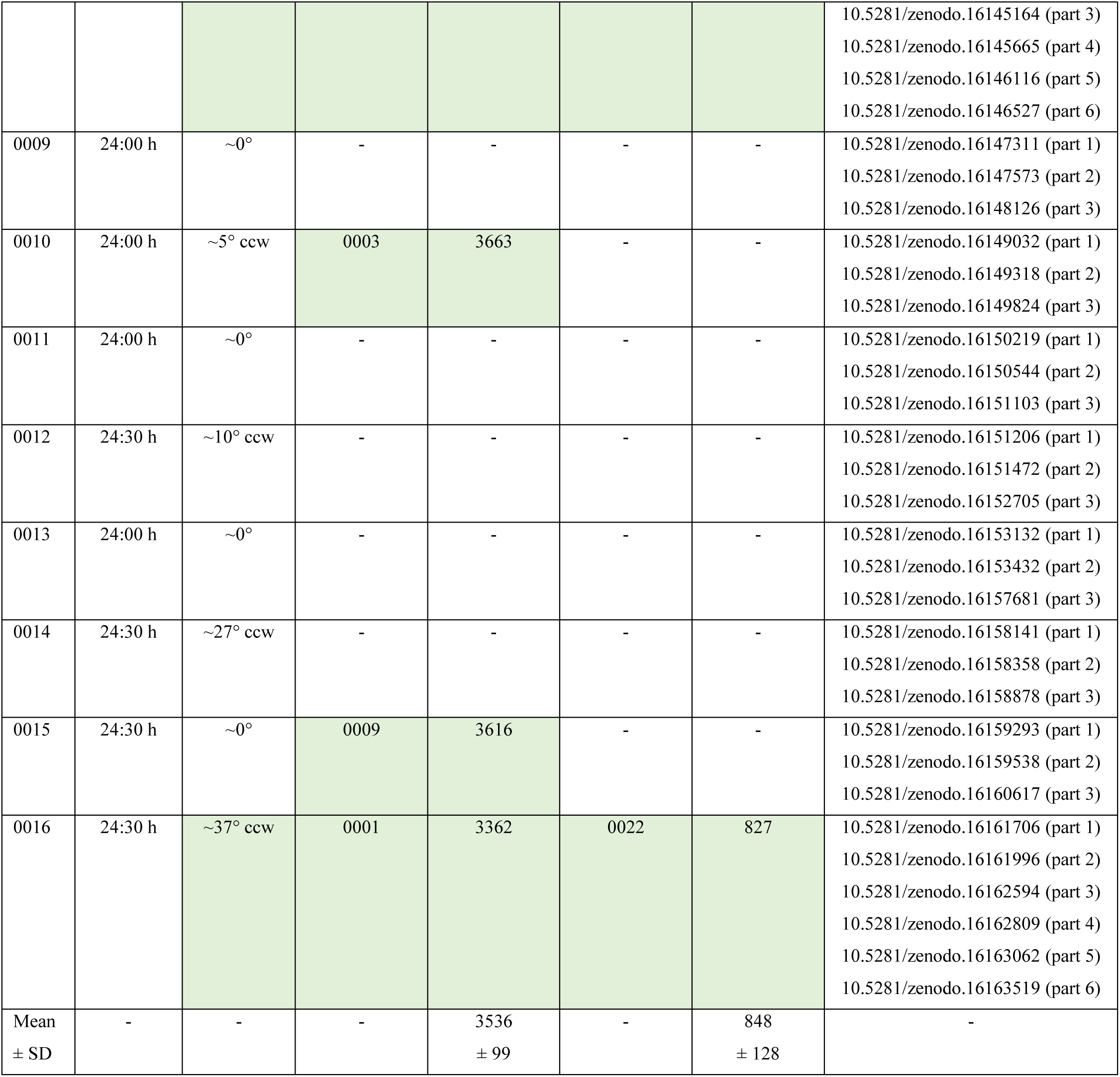
Overview of datasets with selected metadata. Datasets are split into multiple parts for convenient download. The first part contains all files from the basic branch, the second part contains all files from the weighted-average fusion branch and the fusion-deconvolution branch, the third part contains a selection of files from the preliminary branch, and the fourth to sixth parts (optional) contain the preliminary fusion *z* stacks. Green shading in the ‘Embryo rotation’ column indicates datasets in which the embryo rotated by a sufficient angle to enable serosa nuclei segmentation using ‘time-merged’ 3D images; green shading in the ‘Nuclei segmentation’ column indicates the datasets and time points for which nuclei segmentation was performed. DS, dataset; TP, time point; cw, clockwise as seen from the anterior pole; ccw, counter-clockwise as seen from the anterior pole; SD, standard deviation.

The preliminary processing branch (denoted with ‘P’ in subfolder names) encompasses registration and fusion of *z* stacks acquired along multiple directions based on the multiview deconvolution plugin^37^ to obtain fused 3D images with roughly isotropic resolution and diminished shadowing artifacts. Firstly, based on the raw *z* stacks (P1 folder, data only provided for five datasets), *z* and *y* maximum projections were calculated for all *z* stacks and concatenated to time (*t*) stacks (P2 folder). Secondly, the *y* projection *t* stacks were used to estimate the mutual *x*-*z* quadratic footprint of all four *z* stacks recorded along different directions, and the *z* stacks were trimmed accordingly, typically to sizes of 688–780 voxels along *x* and 172–195 voxels (i.e. planes) along *z* (P3 folder). Thirdly, for each direction, the 3D region containing the embryo was almost completely removed (i.e. blacked out) from the trimmed *z* stacks except for a small cap at the top of the 3D image (data not provided). These *ad hoc* stacks were used for registration, i.e. for calculating the transformation matrices based on interest points derived from (i) the fluorescent microsphere ‘cloud’ at the bottom of the image and (ii) a few nuclei in the small cap at the top of the image that were visible from all four directions. Fourthly, the transformation matrices were used to calculate both preliminary weighted-average fusion 3D images as well as preliminary fusion-deconvolution 3D images. The point spread function (PSF) used for deconvolution was derived from the fluorescent microspheres, defined as 23 voxels in all three spatial dimensions, and the operation was run for 10 iterations. Finally, in the raw fused 3D images, the origin (coordinates 0-0-0) was reassigned to the left-upper-front voxel in both fusion derivatives, and the raw fusion-deconvolution 3D images were converted from the resulting 32-bit floating point-based format to a 16-bit integer-based format (P4 and P5 folders).

The resulting images were further processed in the weighted-average fusion branch (denoted with ‘F’ in subfolder names) and in the fusion-deconvolution branch (denoted with ‘D’ in subfolder names) to generate rendering- and segmentation-ready 3D images (Figure 2). Firstly, the fused images were rotated around all three spatial axes so that, at the time point immediately after serosa window closure (cf. Table 1, fifth column), the lateral axes of the embryos aligned with *x*, the anterior-posterior axes with *y* and the ventral-dorsal axes with *z*. Since the anterior-posterior and lateral axes cannot be reliably identified at the uniform blastoderm stage, they cannot be physically aligned with the *x* and *z* axes of the microscope during mounting. Consequently, rotations around *y* reached up to 45°. In contrast, rotations around *x* and *z* were applied to correct minor misalignments with the *y* axis during mounting and were small in magnitude (∼3° on average for both axes). The 3D images were then cropped to a size of 600 × 1000 × 600 voxels, or in the case of large embryos, 600 × 1100 × 600 voxels (F1 and D1 folders). Secondly, based on the axis-aligned and cropped 3D images, *y* and *z* gradient projections (i.e. projections in which front planes contribute more strongly to the final image than the rear planes) were calculated for four directions (in the orientations 0°, 90°, 180° and 270°) and concatenated into *t* stacks (F2 and D2 folders). Thirdly, the axis-aligned and cropped 3D images were concatenated to *t*-*z* hyperstacks, subjected to histogram matching (Bleach Correction → Histogram Matching) to equalize the signal intensity over the entire time course, adjusted in brightness and contrast, and split into 3D images again (F3 and D3 folders). Fourthly, background artifacts were removed by applying two 3D masks along the *x* and *z* axes, respectively (mask region-of-interest files can be found in the B0 folder). Fifthly, based on the axis-aligned, cropped and intensity-adjusted 3D images, *y* and *z* gradient projections were again calculated for four directions (in the orientations 0°, 90°, 180° and 270°) and concatenated into *t* stacks (F4 and D4 folders). Finally, the *t* stacks for all directions were combined into horizontal and vertical *t* stack montages (F5 and D5 folders).

**Figure 2.**
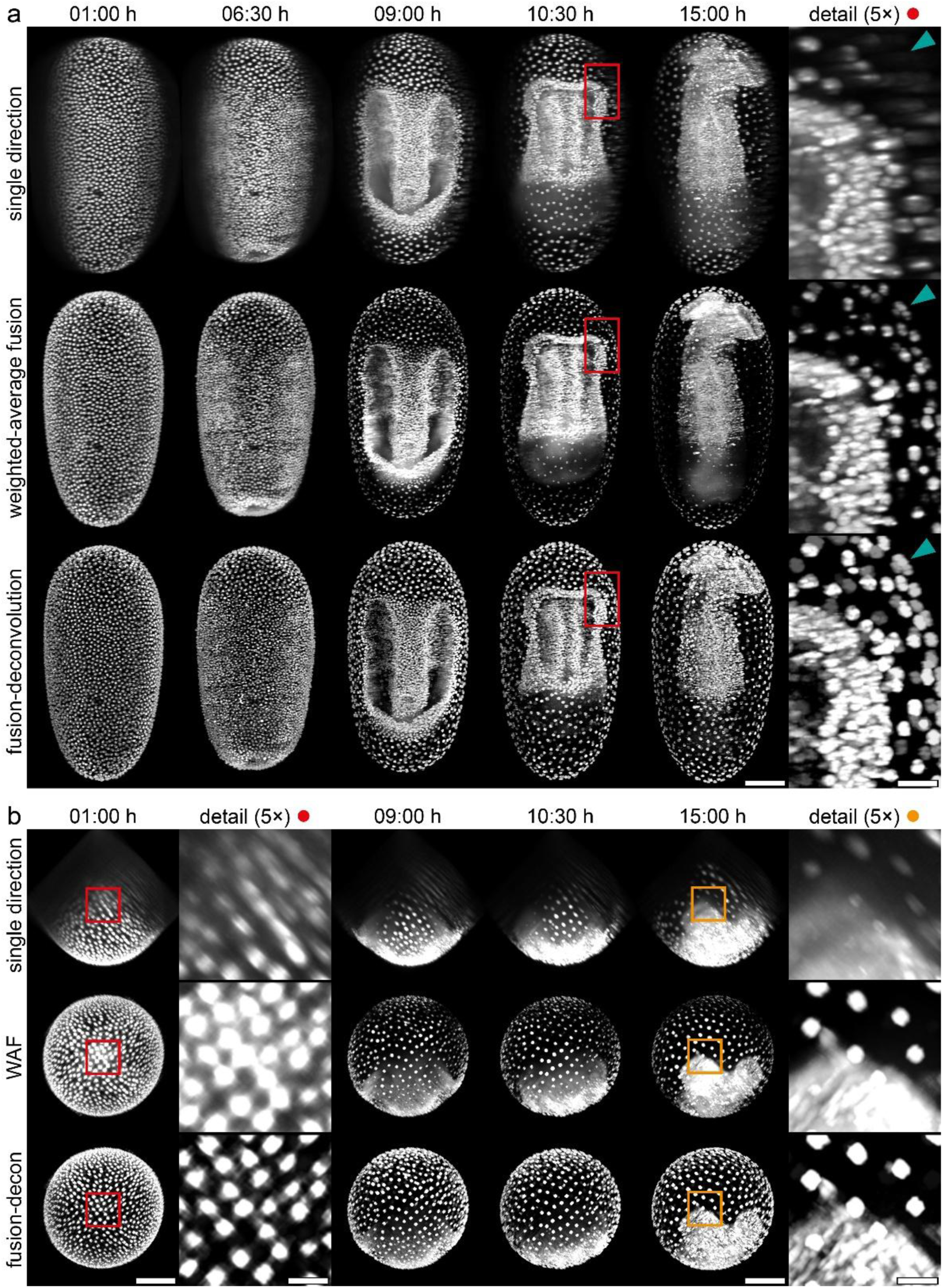
Comparison of 3D images obtained by weighted-average fusion and fusion-deconvolution. **(a)** Juxtaposition of maximum projections along *z* of both *z* stacks and the corresponding fused 3D images at multiple time points. The embryo was initially recorded along four directions in both ventrolateral and both dorsolateral views, and one ventrolateral view was rotated around *y* to generate a ventral view (first row). The four directions were then subjected to both weighted-average fusion and fusion-deconvolution and then rotated around all three spatial axes to align embryonic and image axes (second and third row). Red rectangles indicate the regions shown in the detail images, which display small areas in the anterior ventrolateral region. Serosa cell nuclei in the lateral region are clearly discernible only in the fused representations (teal arrowheads). **(b)** Juxtaposition of maximum projections along *y* of both *z* stacks and the corresponding fused 3D images at multiple time points. Red and orange rectangles indicate the regions shown in the detail images, which display small areas at the anterior pole. Serosa cell nuclei are only weakly recognizable in the non-fused projections due to shadowing artifacts. WAF, weighted-average fusion; fusion-decon, fusion-deconvolution. Image data from dataset DS0005. Scale bars, 100 µm (main images) and 20 µm (detail images).

### Nuclei segmentation

Blastodermal nuclei at the end of blastoderm formation (i.e. the last time point displaying the uniform blastoderm) and serosa nuclei at the onset of germband elongation (i.e. the first time point at which serosa window closure was unequivocally completed) were segmented in 3D images derived from the axis-aligned, cropped, and intensity-adjusted 3D images of the fusion-deconvolution branch (D3 folder). Segmentation was performed manually in the dedicated 3D image visualization and analysis software Arivis Pro (Version 4.2.1, Zeiss Microscopy) using the ‘Magic Wand’ tool in the 2D viewer mode (search tolerance 2–10%, holes not allowed). Since serosa nuclei at the onset of germband elongation were difficult to demarcate due to the proximity of the developing germband, segmentation was restricted to the seven datasets in which the germband rotated by ≥35° around the anterior-posterior axis after serosa window closure (cf. Table 1, third column). In these datasets, serosa nuclei were segmented using ‘time-merged’ 3D images displaying (i) the time point of interest in green and (ii) the final time point of the time series in red. This representation facilitated identification of serosa nuclei based on spatial correspondence across time points (Supplementary Figure 1). In three datasets (DS0003, DS0006, and DS00016), nuclei identification in certain regions was hindered by suboptimal local contrast in the standard intensity-adjusted 3D images. To address this, additional 3D ‘helper’ images with time point-specific brightness and contrast adjustments were generated to support accurate segmentation. Of the seven datasets in which the serosa nuclei were quantified, five also allowed segmentation of blastodermal nuclei (in the remaining two, imaging started after blastoderm rearrangement had already commenced). To match the number of serosa segmentations, blastoderm nuclei were segmented in two further datasets in which the embryos did not undergo substantial germband rotation.

## Data Records

The SLICE-2 datasets are available for download at Zenodo^38^ as ZIP-compressed TIFF files. For each dataset, files from the different image processing branches are distributed across three or six ZIP archives (Table 1). The first archive contains all files from the basic branch (part 1, ‘B’), the second archive contains all files from the weighted-average fusion and fusion-deconvolution branches (part 2, ‘F’ and ‘D’), and the third archive contains selected files from the preliminary branch (part 3, ‘P1’–‘P3’). For the five datasets in which both blastodermal and serosa nuclei were segmented, additional archives provide the preliminary weighted-average fusion 3D images (part 4, ‘P4’) and the preliminary fusion-deconvolution 3D images (part 5 and part 6, ‘P5’). For the remaining eleven datasets, these images can be reproduced using the image data in the P3 folder in combination with the fusion metadata in the B0 folder. Each entry also includes comprehensive experimental metadata in form of a human- and machine-readable XLSX file. An overview of the folder structure is provided in Supplementary Table 1. The complete collection has a total size of 1.65 Terabytes.

## Data Overview

For the segmentation data, a ZIP archive can be downloaded from Zenodo^39^ containing (i) copies of the axis-aligned, cropped, and intensity-adjusted 3D images from the fusion-deconvolution branch (including the time-merged 3D images and ‘helper’ images, if applicable) used for segmentation (Table 1, fourth and fifth column), (ii) the corresponding segmentation SIS files with their associated resources, and (iii) machine-readable nuclei coordinates XLSX files.

## Technical Validation

During the entire project and each experiment, particular emphasis was placed on ensuring dataset homogeneity and comparability. Important technical considerations included:

### Homozygous imaging cultures

After completion of the mating procedure associated with the AGOC vector concept^40^, the AGOC{ATub’H2A/H2B°NB-mEmerald} #1 transgenic subline carries a single homozygous insertion of the transgene^25^. In consequence, the respective imaging culture produced exclusively homozygous embryos, eliminating variation in fluorescence signal strength and/or pattern that may otherwise arise from differences in transgene dosage and/or multiple genomic insertions.

### Microscope calibration and effective laser power

Prior to each recording, the DSLM was calibrated using a two-step routine^33^ to prevent common issues such as offsets or tilts between the light sheet and the focal plane of the detection objective. The effective laser power, defined as the power at the precise location of the embryos during imaging rather than the nominal output of the laser module, was measured regularly. When deviations were detected, the laser output was adjusted to ensure consistent illumination intensity across all dataset acquisitions.

### Mounting arrangement

Since the embryo and the agarose column containing the fluorescent microspheres were positioned along *y* with sufficient distance between them, the illumination and detection light paths to the embryos were not obstructed by the microspheres. In consequence, in all but one dataset (DS0002), the signal from the fluorescent microspheres could be fully removed from the *z* stacks and 3D images by simple horizontal cropping, eliminating the need for additional masking or specialized processing.

### Confirmation of non-invasive imaging

To verify that the imaging procedure (e.g. the light exposure) did not induce developmental defects, all embryos were retrieved after recording, reared to adulthood, and assessed for fertility as described previously^27^. All embryos hatched morphologically intact. Of the resulting larvae, all but the one from DS0009 developed into healthy adults. Among the adults, fertility was confirmed for all individuals except those from DS0004 and DS0010.

### Registration quality

Registration quality was evaluated for each dataset to ensure accurate alignment of the *z* stacks. Two quantitative metrics were assessed: the residual registration error and the interest-point correspondence ratio. Across all datasets, the residual error remained typically below one pixel, indicating high spatial fidelity of the registration. The correspondence ratio consistently exceeded >90%, reflecting reliable identification and alignment of landmarks (i.e. fluorescent microspheres) across *z* stacks acquired along multiple directions (Supplementary Figure 2).

### Validation of image registration and fusion

Correct image registration and fusion were verified through manual visual inspection of *ad hoc* projections of the raw fused 3D images that still retain signal from the fluorescent microspheres, with particular attention to the microsphere-derived signal, which appears compass-star-shaped in *y* projections of correctly fused reconstructions (Figure 3). Across a total of 786 independent fusion processes, only a single error was identified. In DS0010, the four *z* stacks of TP0013 were incorrectly fused in the initial run (Supplementary Figure 3) as well as in two subsequent repetitions. As a workaround, fusion was performed using the transformation matrices of TP0014. This incident has been documented in the corresponding metadata.

**Figure 3.**
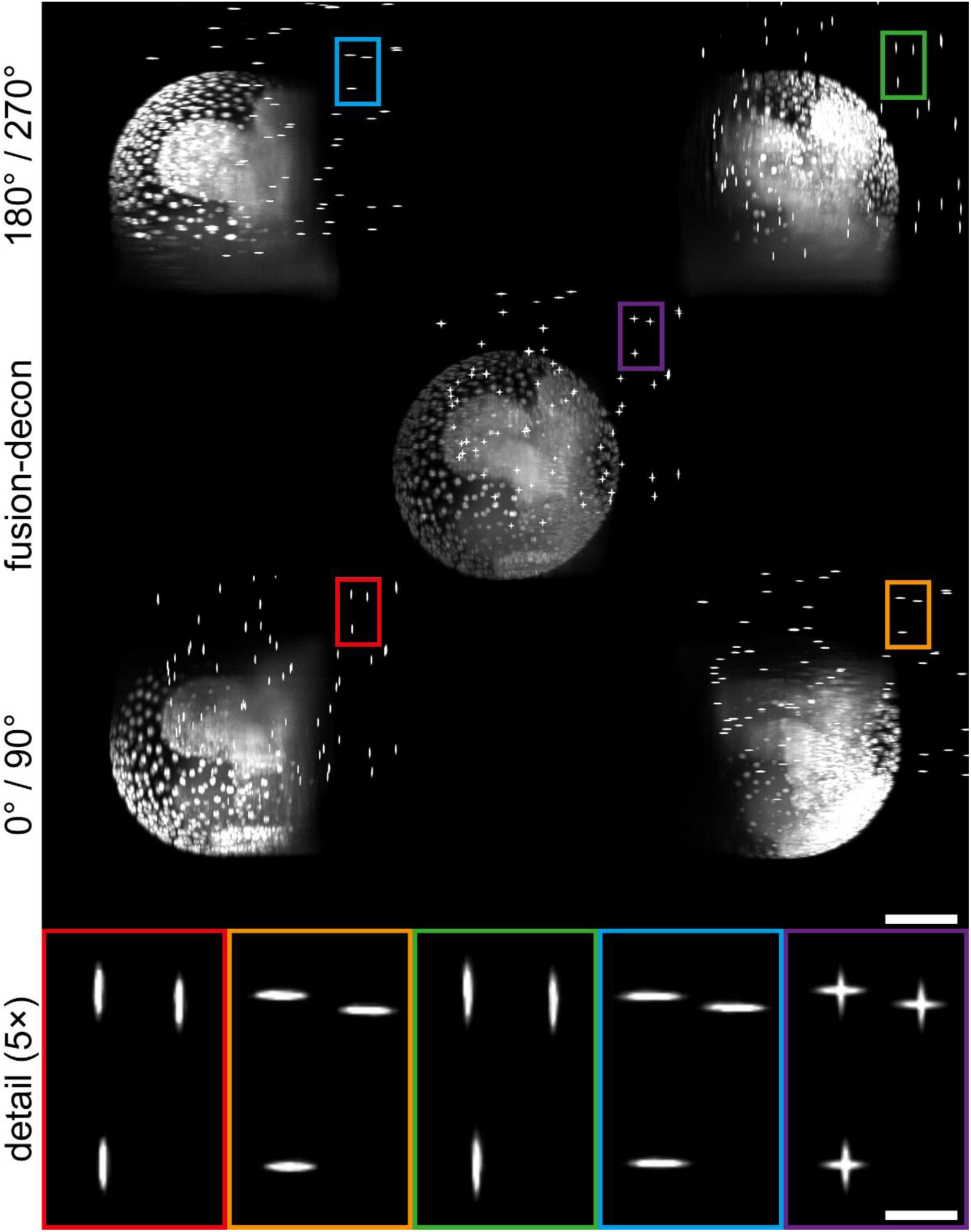
Fluorescent microsphere-based validation of registration and fusion. Maximum projections along *y* of *ad hoc z* stacks that were trimmed to a mutual quadratic footprint (outer panels) and the corresponding fused 3D image prior to rotation and cropping, i.e. with the signal from the fluorescent microsphere cloud still present (inner panel). All five detail images show the same three fluorescent microspheres. The compass-star-like geometry of the signal in the detail image derived from the fused 3D image indicates correct fusion. Notably, the 3D image-derived projection exhibits a smaller point spread than the individual *ad hoc z* stack projections, reflecting the effect of deconvolution. Fusion-decon, fusion-deconvolution. Image data from dataset DS0005. Scale bars, 100 µm (main images) and 20 µm (detail images).

### Manual segmentation by two researchers

Manual segmentation of blastodermal nuclei at the end of blastoderm formation as well as serosa nuclei at the beginning of germband elongation was performed by two researchers to ensure accuracy and reproducibility. Following the initial segmentation by the first researcher, all segmented nuclei were systematically reviewed by the second researcher. The review was carried out using both the 2D viewer and the 3D viewer modes in Arivis Pro. This double-checking procedure was implemented to minimize user bias and segmentation errors.

## Usage Notes

The image data structure (cf. Data Records section) is compatible with any software capable of handling 3D dynamic image data. Fiji^35^, an open-source framework widely used for working with multi-dimensional microscopy data, is recommended as the primary platform for image processing and analysis.

Regarding the segmentation data, the SIS files and their accompanying resources can be opened with Arivis Pro, Version 4.2.1 or higher (Figure 4A). The nuclei coordinates CSV files can be used independently or in conjunction with respective image data TIFF files e.g. to generate overlays. This can be done, for instance, with the 3D Viewer plugin^41^ for Fiji, which comes with a CSV import function (Figure 4B). In addition, the Dash web application framework for Python^42^ enables browser-based 3D visualization of nuclei positions with arbitrary rotation (Figure 4C, a respective script is provided, cf. Code Availability section).

**Figure 4.**
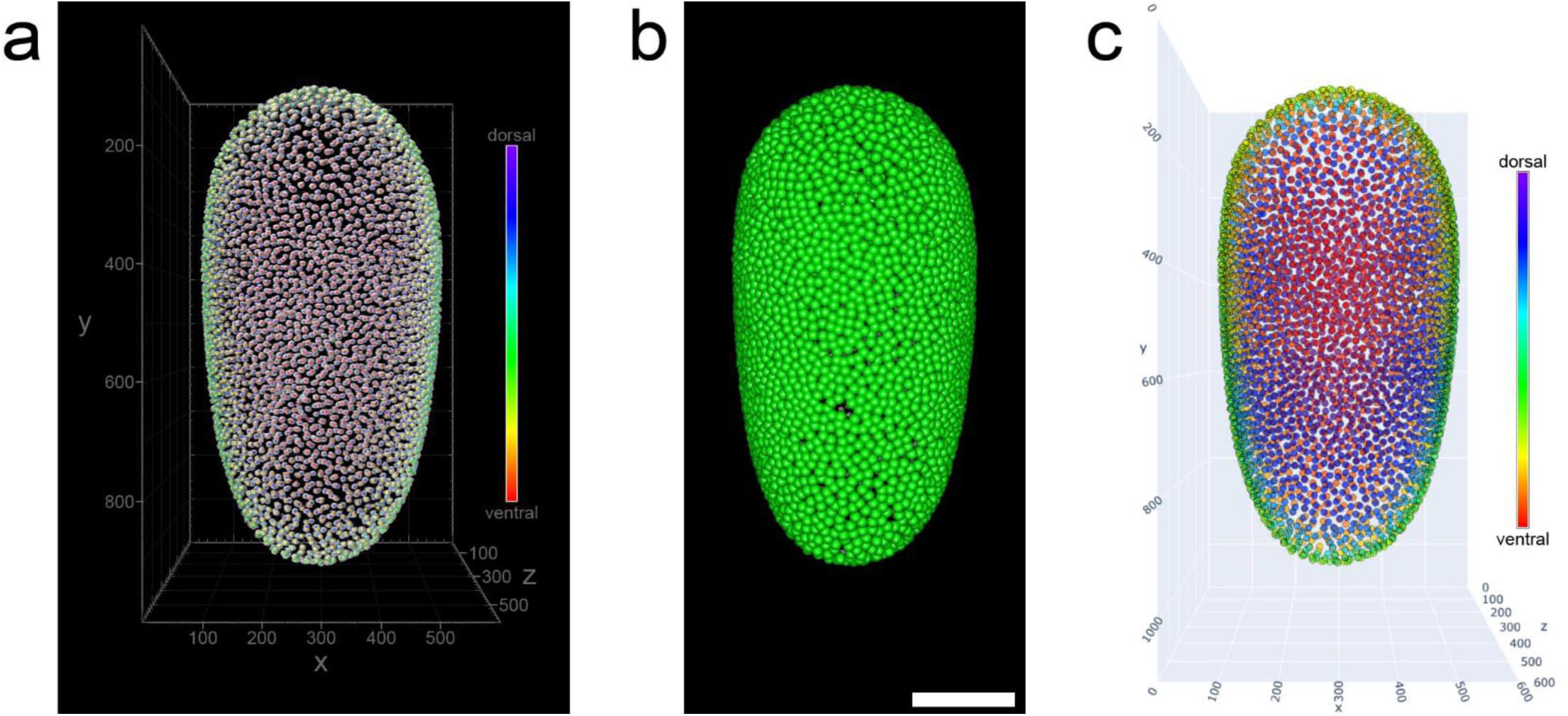
Illustration of nuclei segmentation using three visualization programs. **(a)** Transparency-based volume rendering using Arivis Pro. This program enables overlay visualization of the voxel data (gray-transparent) and the centroids of segmented nuclei (ventral-dorsal heatmap). The bounding box indicates voxel dimensions. **(b)** Volume rendering using the 3D Viewer plugin in Fiji. This program also supports overlay visualization of voxel data and centroid positions (green), but the absence of a transparency mode requires increasing the diameter of the centroid marker spheres; otherwise, the markers disappear within the nuclei. Heatmap coloring is not supported. Scale bar, 200 pixels. **(c)** Volumetric reconstruction of centroid positions of the segmented nuclei (ventral-dorsal heatmap) using the Nuclei Pocket Viewer 3D Python script. This program does not support voxel data rendering. The bounding box indicates voxel dimensions. Image data from dataset DS0005.

## Data Availability

The Zenodo link^38^ redirects to the SLICE-2 dataset collection overview page, which allows entries to be sorted by various convenient criteria. Entries belonging to the same dataset are cross-linked with their respective other parts in the ‘Additional details’ section. Digital Object Identifiers (DOIs) for all entries are provided in Table 1 and can be resolved via the DOI website (https://www.doi.org). An additional entry on Zenodo allows independent download of the segmentation data^39^.

## Code Availability

All Fiji macros used for image processing are available on GitHub (https://github.com/BugCube/FijiFusionMacros). They are numbered according to the processing workflow and additionally tagged to reflect the subfolder structure (cf. Methods section). Macros labeled ‘Universal’ require no user input, those labeled ‘Parameters’ require user-defined input, and those labeled ‘ParametersDefault’ accept optional input, with default values provided. The Dash-based Python script for web browser-based 3D visualization of nuclei positions, ‘Nuclei Pocket Viewer 3D’, is also available on GitHub (NucleiPocketViewer3D_V1.0.py, https://github.com/BugCube/NucleiPocketViewer3D).

## Acknowledgements

We thank Ernst H.K. Stelzer for his generous support, including resources and scientific guidance, and Sven Plath for technical assistance.

## Author Contributions

FK and FS conceived the research, performed the imaging experiments, and processed the data. FS prepared the display items and wrote the manuscript with input from FK and SM.

## Supplementary Material

**Supplementary Figure 1.**
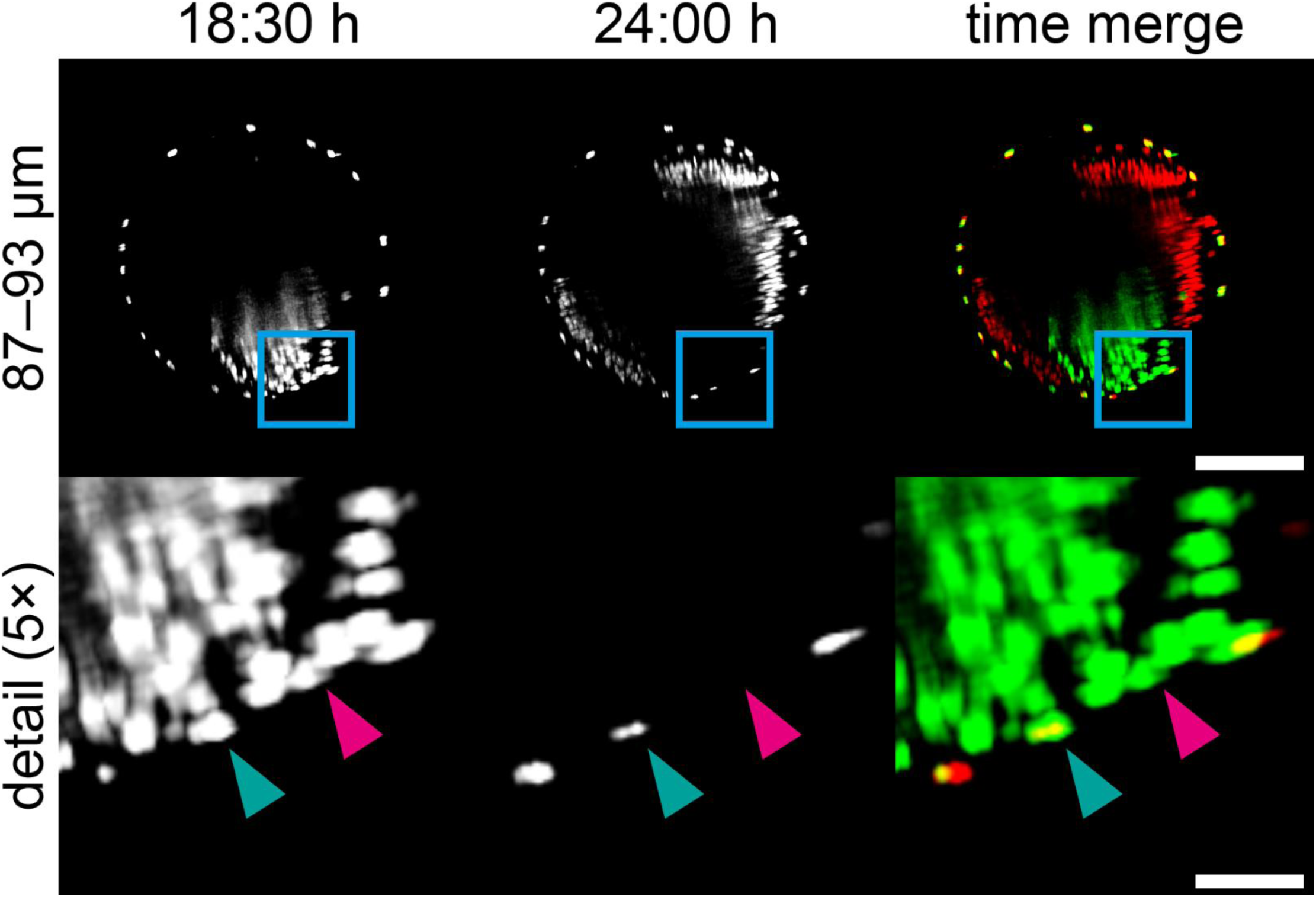
Identification of serosa nuclei in close proximity to the developing germband using ‘time-merged’ 3D images. Short-distance (∼6 µm) maximum projections along *y* of a ‘time-merged’ 3D image. Between the stage immediately after serosa window closure (first column), and the last time point of the dataset (second column), the germ band rotates ∼48° clockwise (as viewed from the anterior pole) around the anterior-posterior axis (i.e. the *y* axis) within the serosa. This rotation enables identification of serosa nuclei in the time-merge volume: nuclei that belong to the serosa retain signal at the same spatial position across both time points, whereas nuclei that belong to the amnion/germband do not (third column). Blue rectangles indicate the regions shown in the detail images, which display a small area where the serosa is in contact with the germband at 18:30 h but no longer at 24:00 h. Based on the presence or absence of overlapping signal across the two time points, the nucleus marked by the teal arrowhead was classified as belonging to the serosa, while the nucleus marked by the pink arrowhead was not. Image data from dataset DS0002. Scale bars, 100 µm (main images) and 20 µm (detail images).

**Supplementary Figure 2.**
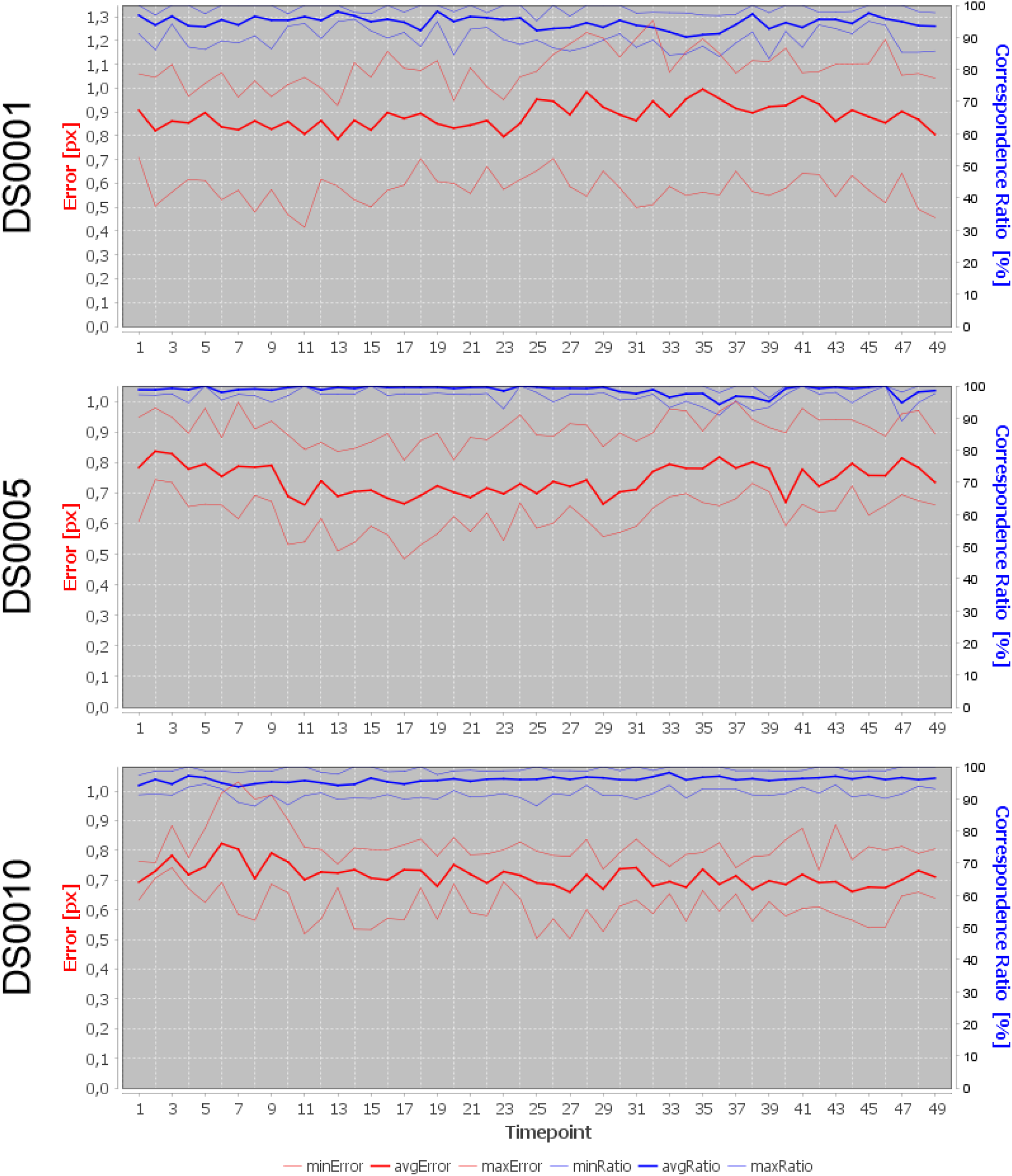
Registration quality assessment. The graphs summarize multiview registration performance across the entire acquisition period for three representative datasets. Shown are the minimum, average, and maximum registration error in pixels (minError, avgError, maxError; red curves, left axis), and the minimum, average, and maximum correspondence ratio (minRatio, avgRatio, maxRatio; blue curves, right axis) computed across the six direction pairs (1↔2, 1↔3, 1↔4, 2↔3, 2↔4, and 3↔4). The correspondence ratios indicate the fraction of identified fluorescent microspheres that were successfully matched across the four acquisition directions, i.e. how many correspondences the registration algorithm could establish. Across all three datasets, the average registration error remains below 1 pixel, which is small relative to the final dimensions of the fused 3D image (i.e. 600 × 1000 × 600 voxels). At the same time, the correspondence ratios consistently exceed 90%, showing that the vast majority of fluorescent microspheres were successfully matched in all direction pairs. Together, these metrics demonstrate that the multiview registration is reliable and robust, providing a strong basis for high-quality fusion and downstream quantitative analyses.

**Supplementary Figure 3.**
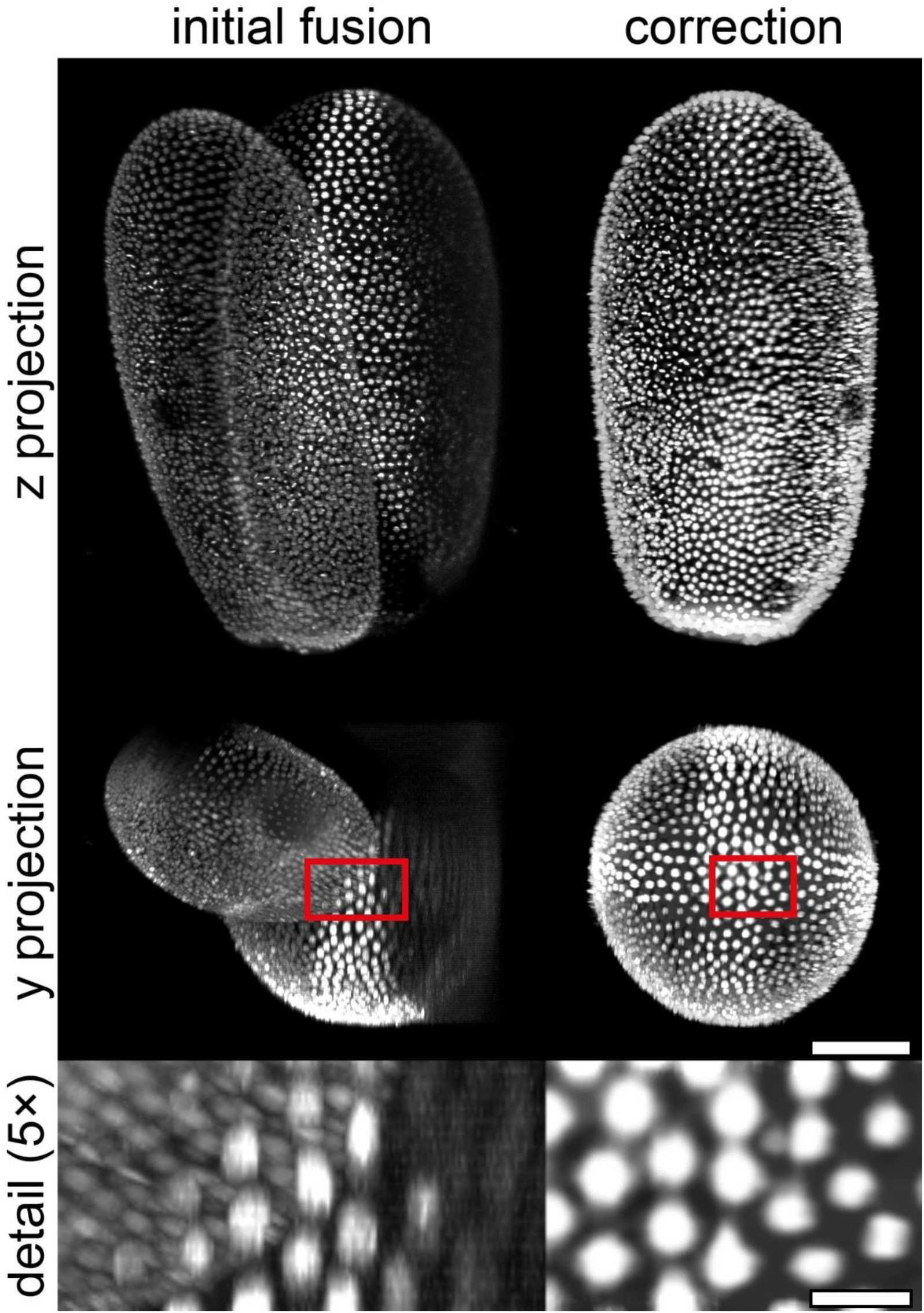
Fusion error in DS0010. At the 13^th^ timepoint (TP0013), multiple *z* stacks were incorrectly positioned during the fusion process. After two repetitions with slightly altered parameters produced similar issues, the fusion was repeated using the registration parameters from the subsequent time point. This was the only such error across all time points of the sixteen datasets. Red rectangles indicate the regions shown in the detail images, which display an area at the anterior pole. In the initial fusion, the nuclei located directly at the pole were likely contributed by only a single acquisition direction. Image data from dataset DS0010. Scale bars, 100 µm (main images) and 20 µm (detail images).

**Supplementary Table 1.** Data folder structure overview for the image processing branches. The folders are grouped into four branches: the basic branch (‘B’), the fusion-deconvolution branch (‘D’), the weighted-average fusion branch (‘F’), and the preliminary processing branch (‘P’). In the folder names, ‘N’ serves as a placeholder that indicating the type of brightness/contrast processing applied: either histogram correction followed by intensity adjustment (abbreviated as ‘HS’ in *z* stacks file names or ‘ZH’ in *z* maximum projection file names), or only intensity adjustment (abbreviated as ‘AS’ in *z* stacks file names or ‘ZA’ in *z* maximum projection file names). Bold formatting highlights quick-start information, while italics denote explanatory details related to the subfolder hierarchy. RelVol, approximate estimate of relative data volume within a dataset.

| Branch | Folder | RelVol | Rationale |
| --- | --- | --- | --- |
| B branch | (B0)-Metadata | - | Contains fusion metadata. The comprehensive dataset metadata XLSX file is normally stored in this folder but has been extracted to facilitate independent access without downloading the full dataset. |
|  | (B1)-ZStacks-ZS | ++ | Derives from the (P1) folder. Contains non-fused rotated and cropped *z* stacks ('ZS'). *The folder contains subfolder layers for channels and directions.* |
|  | (B2)-TStacks-ZM | + | Contains *t* stacks ('TS') of *z* maximum projections ('ZM') derived from the non-fused, rotated, and cropped *z* stacks. *The folder contains subfolders for channels and directions.* |
|  | (B4)-TStacks-ZN | + | Contains *t* stacks ('TS') of *z* maximum projections ('ZH') derived from the non-fused, rotated, and cropped *z* stacks that were subjected to histogram matching and intensity adjustment. *The folder contains subfolders for channels and directions.* |
|  | (B5)-TStacks-ZN-Montage | + | Contains horizontal and vertical direction montages ('AX' and 'AY', respectively) of *t* stacks ('TS') derived from the *z* maximum projections ('ZH') of non-fused, rotated, and cropped *z* stacks that were subjected to histogram matching and intensity adjustment. **The montages are intended for quick visual inspection of the respective dataset/processing branch.** |
| D branch | (D1)-ZStacks-Decon-ZS | +++ | Derives from the (P5) folder. Contains fusion-deconvolution-derived, rotated, and cropped 3D images ('ZS'). *The folder contains two subfolder layers for channels and directions.* |
|  | (D2)-TStacks-Decon-YM-ZM | + | Contains *t* stacks ('TS') of *y* and *z* maximum projections ('YM' and 'ZM', respectively) derived from the fusion-deconvolution-derived, rotated, and cropped 3D images. *The folder contains two subfolder layers for channels and directions.* |
|  | (D3)-ZStacks-Decon-NS | + | Contains fusion-deconvolution-derived, rotated, and cropped *z* stacks ('NS') that were subjected to histogram matching and intensity adjustment. **Segmentation data derive from the files in this folder.** *The folder contains two subfolder layers for channels and directions.* |
|  | (D4)-TStacks-Decon-YN-ZN | + | Contains *t* stacks ('TS') of *y* and *z* maximum projections ('YH' and 'ZH', respectively) derived from the fusion-deconvolution-derived, rotated, and cropped 3D images that were subjected to histogram matching and intensity adjustment. *The folder contains two subfolder layers for channels and directions.* |
|  | (D5)-TStacks-Decon-ZN-Montage | + | Contains horizontal and vertical direction montages ('AX' and 'AY', respectively) of *t* stacks ('TS') derived from the *z* maximum projections ('ZH') of fusion-deconvolution-derived, rotated, and cropped *z* stacks that were subjected to histogram matching and intensity adjustment. **The montages are intended for quick visual inspection of the respective dataset/processing branch.** |
| F branch | (F1)-ZStacks-Fusion-ZS | ++ | Derives from the (P4) folder. Contains weighted-average fusion-derived, rotated, and cropped 3D images ('ZS'). *The folder contains two subfolder layers for channels and directions.* |
|  | (F2)-TStacks-Fusion-YM-ZM | + | Contains *t* stacks ('TS') of *y* and *z* maximum projections ('YM' and 'ZM', respectively) derived from the weighted-average fusion-derived, rotated, and cropped 3D images. *The folder contains two subfolder layers for channels and directions.* |
|  | (F3)-ZStacks-Fusion-NS | + | Contains weighted-average fusion-derived, rotated, and cropped *z* stacks ('NS') that were subjected to histogram matching and intensity adjustment. *The folder contains two subfolder layers for channels and directions.* |
|  | (F4)-TStacks-Fusion-YN-ZN | + | Contains *t* stacks ('TS') of *y* and *z* maximum projections ('YM' and 'ZM', respectively) derived from the weighted-average fusion-derived, rotated, and cropped 3D images that were subjected to histogram matching and intensity adjustment. *The folder contains two subfolder layers for channels and directions.* |
|  | (F5)-TStacks-Fusion-ZN-Montage | + | Contains horizontal and vertical direction montages ('AX' and 'AY', respectively) of *t* stacks ('TS') derived from the *z* maximum projections ('ZH') of weighted-average fusion-derived, rotated, and cropped *z* stacks that were subjected to histogram matching and intensity adjustment. **The montages are intended for quick visual inspection of the respective dataset/processing branch.** |
| P branch | (P1)-ZStacks-RS | ++++ | Contains raw *z* stacks ('RS'). Feeds into the B branch. **This folder contains the top-level data in the processing hierarchy.** |
|  | (P2)-TStacks-YR-ZR | + | Contains *t* stacks ('TS') of *y* and *z* maximum projections derived from the raw *z* stacks ('YR' and 'ZR', respectively). |
|  | (P3)-QStacks | +++ | Contains *z* stacks cropped to a quadratic footprint ('QS'). |
|  | (P4)-QStacks-Fusion | +++ | Contains the preliminary weighted-average fusion 3D images. Data only provided for the five datasets in which both blastodermal and serosa nuclei were segmented. Feeds into the F branch. |
|  | (P5)-QStacks-Decon | ++++ | Contains the preliminary fusion-deconvolution 3D images. Data only provided for the five datasets in which both blastodermal and serosa nuclei were segmented. Feeds into the D branch. |

**Supplementary Movie 1 – Representative blastodermal nuclei segmentation.** Transparency-based volume rendering of a fused-deconvolved 3D image of a *Tribolium castaneum* embryo at the uniform blastoderm stage, showing the voxel data in gray-transparent and overlaid with the centroids of the segmented superficial blastodermal nuclei as a ventral-dorsal heatmap. In the first part of the movie, the embryo is rotated around the *y* axis; in the second part, the embryo is rotated around the *x* axis; and in the third part, the embryo is shown from a ventral–anterior oblique perspective. The grid spacing of the bounding box is 100 pixels. Image data from dataset DS0005. Frame rate is 30 frames per second.

**Supplementary Movie 2 – Representative serosa nuclei segmentation.** Transparency-based volume rendering of a fused-deconvolved 3D image of a *Tribolium castaneum* embryo at the beginning of germband elongation, showing the voxel data in gray-transparent and overlaid with the centroids of the segmented serosa nuclei as a ventral-dorsal heatmap. In the first part of the movie, the embryo is rotated around the *y* axis; in the second part, the embryo is rotated around the *x* axis; and in the third part, the embryo is shown from a ventral–anterior oblique perspective. The grid spacing of the bounding box is 100 pixels. Image data from dataset DS0005. Frame rate is 30 frames per second.

